# Microglia Regulate Neuroglia Remodeling in Various Ocular and Retinal Injuries

**DOI:** 10.1101/366310

**Authors:** Eleftherios I. Paschalis, Dylan Lei, Chengxin Zhou, Xiaohong Nancy Chen, Vassiliki Kapoulea, Pui-Chuen Hui, Reza Dana, James Chodosh, Demetrios Vavvas, Claes H. Dohlman

## Abstract

Reactive microglia and infiltrating peripheral monocytes have been implicated in many neurodegenerative diseases of the retina and central nervous system (CNS). However, their specific contribution in retinal degeneration remains unclear. We recently showed that peripheral monocytes that infiltrate the retina after ocular injury in mice become permanently engrafted into the tissue, establishing a pro-inflammatory phenotype that promotes neurodegeneration. Here, we show in mice that microglia regulate the process of neuroglia remodeling during ocular injury, and their depletion results in marked upregulation of inflammatory markers, such as *Il17f*, *Tnfsf11*, *Ccl4*, *Il1a*, *Ccr2*, *Il4*, *Il5*, and *Csf2* in the retina, abnormal engraftment of peripheral CCR2^+^ CX3CR1^+^ monocytes into the retina and is associated with increased retinal ganglion cell (RGC) loss, retinal nerve fiber layer thinning, and RPE65^+^ cell migration onto the retinal surface. Furthermore, we show that other types of ocular injuries, such as penetrating corneal trauma and ocular hypertension, also cause similar changes. However, optic nerve crush injury mediated RGC loss evokes neither peripheral monocyte response in the retina, nor RPE65^+^ cell migration, although peripheral CX3CR1^+^ and CCR2^+^ monocytes infiltrate the optic nerve injury site and remain present for months. Our study suggests that microglia are key regulators of peripheral monocyte infiltration and RPE migration and their depletion results in abnormal neuroglia remodeling that exacerbates neuroretinal tissue damage. This mechanism of retinal damage through neuroglia remodeling may be clinically important for the treatment of patients with ocular injuries, including surgical traumas.

## Introduction

Microglia are specialized phagocytes of the retina and central nervous system (CNS) (1–3) responsible for the elimination of noxious stimuli, either by phagocytosis or cytokine release (4–11). Reactive microglia have been identified in almost all neuroretinal diseases and injuries (12), however, their exact role is not yet fully elucidated. Upon activation, microglia have been shown to secrete neurotrophic factors that promote neuronal survival and regeneration (4, 13–15) or release inflammatory cytokines that cause cell death (16–19). This functional duality has been confirmed by single-cell RNAseq techniques where microglia were shown to have a heterogeneous transcriptional profile, with signatures ranging from inflammatory to neuroprotective (10, 20). The mechanism by which this phenotype is selected is not understood, yet growing evidence suggests that inflammatory molecules may play a significant role in epigenetic reprogramming of these cells and their phenotype (21–23). Currently, there is no consensus regarding their specific role in ocular diseases and injuries, which appears to be context dependent. Therefore, elucidating the role of microglia can be important for the development of therapies for patients with neuroinflammatory disorders.

We recently showed that microglia may not be the direct mediators of retinal inflammation after acute ocular surface injury. Instead, peripheral monocytes infiltrate into the retina within hours, initiating neuroretinal inflammation and microglia activation (19). Interestingly, peripheral monocytes that infiltrate into the retina after injury do not vanish from the tissue over time. Instead, they engraft permanently, migrating into the three distinct microglia strata (ganglion cell layer: GCL, inner plexiform/inner nuclear layer: IPL/INL, outer plexiform layer: OPL), assume a ramified morphology, and become morphologically indistinguishable from tissue resident microglia (24). However, these cells remain phenotypically different despite their quiescent morphology and engraftment into the retina, retaining MHC-II expression for the long-term. This process of peripheral monocyte engraftment is regulated by yolk sac-derived microglia, and microglia depletion leads to subsequent repopulation by peripheral CX3CR1^+^ monocytes (24). Considering that microglia and macrophages express common cell markers, distinguishing between the two cell populations is not easy, and often requires novel fate mapping techniques (5, 19, 25–27). Instead, by using busulfan myelodepletion and bone marrow CX3CR1^+/EGFP^::CCR2^+/RFP^ chimeras, we showed that peripherally engrafted monocytes remain transcriptionally different compared to microglia (24). This remodeled neuroglia is characterized by new peripherally-derived microglia that are resistant to depletion by CSF1R inhibitor and continue to phagocytose neuroretinal tissue long after the noxious stimuli is removed. We hypothesize that the observed duality in the role of microglia may be a result of a mixed population of peripheral and non-peripheral microglia with diverse progenies as previously suggested (24).

In this study, we used our CX3CR1^+/EGFP^::CCR2^+/RFP^ chimera model (28) and CSF1R inhibitor for microglia depletion to investigate the role of peripheral monocytes and microglia in retinal degeneration and inflammatory expression after various types of ocular injuries.

## Materials and Methods

### Mouse models

All animal-based procedures were performed in accordance with the Association For Research in Vision and Ophthalmology Statement for the Use of Animals in Ophthalmic and Vision Research, and the National Institutes of Health Guidance for the Care and Use of Laboratory Animals. This study was approved by the Animal Care Committee of the Massachusetts Eye and Ear Infirmary. B6.129P-*Cx3cr1^tm1Litt^*/J (CX3CR1^EGFP/EGFP^ Stock 005582) (29), B6.129(Cg)-*Ccr2^tm2.1lfc^*/J (CCR2^RFP/RFP^ Stock 017586) (30), B6.Cg-Tg(*Thy1-YFP*)16Jrs/J (Thy1^*YFP/YFP*^ Stock 003709) (31) mice were obtained from the Jackson Laboratory (Bar Harbor, ME, USA). Mice were bred and used at the ages of 6-12 weeks.

### Ocular hypertension injury

Anesthetized mice were mounted in a stereotaxic frame (39463001; Leica, Buffalo Grove, IL) with Cunningham mouse adaptor (39462950; Leica). The head was immobilized, and a 30 gauge beveled needle connected to a sterile container with balance salt solution plus (BSS Plus,) was introduced into the anterior chamber through clear cornea incision, adjacent to the limbus. The BSS container was elevated to create hydrostatic pressure of 100 mmHg into the eye and this pressure was maintained for 45 minutes. At the completion of the ocular hypertension (OHT) experiment, the needle was carefully removed from the eye and treated with a drop of topical Polytrim^®^ antibiotic (polymyxin B/trimethoprim, Bausch & Lomb Inc, Bridgewater, NJ, USA). The eye was monitored to ensure that the wound has self-sealed and that was not hypotonous.

### Penetrating corneal injury

Penetrating corneal injury was performed by placing a single 11-0 nylon suture (2881G: 11-0 Ethilon black 5″ BV50-3 taper, ©Johnson & Johnson Medical N.V., Belgium) into the cornea of anesthetized mice. A drop of topical Polytrim^®^ antibiotic was applied after the procedure.

### Optic nerve crush injury model

Anesthetized mice were placed under the microscope and a small incision was performed in the temporal conjunctiva. The optic nerve was gently exposed intraorbitally and crushed 0.5-1mm posteriorly to the globe using Dumont #5/45 forceps (F.S.T., #11251-35) for 5 s, while avoiding damage to the adjacent retinal artery. Immediately after the optic nerve crush, the eye was treated with bacitracin ophthalmic ointment (NDC 0574-4022-35, Perrigo) and surgical lubricant (NDC 17238-610-15, HUB Pharmaceuticals. LLC.). Mice were placed on a heating pad and monitored until fully recovered. For analysis, 2 months optic nerve crush mice were euthanized, eyes were removed with more than 5mm of optic nerve attached, and fixated in PFA 4% for 1 hour. The optic nerve was then dissected from the eye at the sclera level and mounted horizontally on a glass slide for confocal microscopy.

### Corneal chemical injury

Alkali chemical burns were performed according to our previous study.(32) In brief, mice were anesthetized using ketamine (60 mg/kg) and xylazine (6 mg/kg) and deep anesthesia was confirmed by a toe pinch test. Proparacaine hydrocloride USP 0.5% (Bausch and Lomb, Tampa, FL, USA) eye drop was applied to the cornea, and after 1 minute was carefully dried with a Weck-Cel (Beaver Visitec International, Inc, Waltham, MA, USA). A 2 mm diameter filter paper was soaked in 1 M sodium hydroxide (NaOH) solution for 10 seconds, dried of excess NaOH by a single touch on a paper towel and applied onto the mouse cornea for 20 seconds. Complete adherence of the filter paper on the corneal surface was confirmed by gentle push of the perimeter using forceps. After the filter paper was removed, prompt irrigation with sterile saline for 10 seconds was applied using a 50 ml syringe with a 25G needle. The mouse was then placed on a heating pad, positioned laterally, and the eye irrigated for another 15 minutes at low pressure using sterile saline. Buprenorphine hydrochloride (0.05 mg/kg) (Buprenex Injectable, Reckitt Benckiser Healthcare Ltd, United Kingdom) was administered subcutaneously for pain management. A single drop of topical Polytrim^®^ antibiotic was administered after the irrigation. Mice were kept on the heating pad until fully awake.

### Tissue preparation for flat mount imaging and RPE65 staining

Following the alkali burn, eyes were enucleated at predetermined time points and fixed in 4% paraformaldehyde (PFA) (Sigma-Aldrich, St Louis, MO, USA) solution for 1 hour at 4ºC. The cornea and retina tissue were carefully isolated using microsurgical techniques and washed 3 times for 5 minutes in phosphate buffer solution (PBS) (Sigma-Aldrich, St Louis, MO, USA) at 4ºC. The tissues were then blocked using 5% bovine serum albumin (Sigma-Aldrich, St Louis, MO) and permeabilized using 0.3% Triton-X (Sigma-Aldrich, St Louis, MO, USA) for 1 hour at 4ºC. The specific antibody was added in blocking solution, incubated overnight at 4ºC, and then washed 3 times for 10 minutes with 0.1% Triton-X in PBS. Tissues were transferred from the tube to positive charged glass slides. Four relaxing incisions from the center to the periphery were made to generate 4 flat tissue quadrants. VECTRASHIELD^®^ mounting medium (Vector Laboratories, H-1000, Vector Laboratories, CA, USA) was placed over the tissue followed by a coverslip. Monoclonal antibody against RPE65 (E-5: sc-390787, Santa Cruz Biotechnology, Inc, Dallas, TX, USA) was used per manufacturer’s protocol, and RPE65^+^ cells were imaged using an epifluorescent microscope (Zeiss Axio Imager M2, Zeiss, Germany).

### TUNEL labeling and quantitation of DNA fragmentation

Following the alkali burn, eyes were enucleated and cryosectioned. TUNEL labeling was performed using a Roche TUNEL kit (12156792910), (Roche, Indianapolis, IN, USA) as previously published.^(33)^ Mounting medium with DAPI (UltraCruz™, Santa Cruz Biotechnology, sc-24941, Dallas, TX, USA) was placed over the tissue followed by a coverslip. Images were taken with an epifluorescent microscope (Zeiss Axio Imager M2, Zeiss, Germany), using the tile technique. DAPI signal (blue) was overlayed with Texas red (TUNEL^+^ cells) and quantified using ImageJ version 1.5s (http://imagej.nih.gov/ij/; provided in the public domain by the National Institutes of Health, Bethesda, MD, USA) to measure the number of TUNEL^+^ cells overlapping with DAPI in the areas of interest. All experiments were performed in triplicate.

### Retinal flat mount imaging

Eyes were enucleated together with their optic nerves at different time points. Retinal and optic nerve tissues were prepared for flat mount using a confocal microscope (Leica SP-5, Leica Microsystems inc, Buffalo Grove, IL, USA). Images were taken at 10x, 20x, 40x and 63x objective lenses using z-stack of 0.7, 0.6, 0.4 and 0.3 μm step, respectively. Image processing with ImageJ version 1.5s was used to obtain maximum and average projections of the z-axis, color depth maps, and 3-D volumetric images. Retinal microglia/macrophage cells were quantified by layer-by-layer technique, total z-stack projection technique, and volumetric 3-D analysis. Retinal nerve fiber layer was quantified by using z-stack projection and binary image conversion or by orthogonal transverse 2-D cuts of the nerves and area measurements.

### Bone marrow chimerism

C57BL/6J mice were myelodepleted with 3 intraperitoneal injections of busulfan (Sigma-Aldrich, St Louis, MO, USA) (35mg/kg) at 7, 5, and 3 days prior to bone marrow transfer. CX3CR1^+/EGFP^::CCR2^+/RFP^, B6::CX3CR1^+/EGFP^ and CX3CR1^CreERT2^::ROSA26-tdTomato bone marrow cells (5×106 total bone marrow cells) were transferred to the myelodepleted C57BL/6J mice through tail vein injection 1 month prior to corneal alkali burn. Bactrim (trimethoprim/sulfamethoxazole 80 mg/ 400 mg, respectively, was resuspended in 400mL drinking water) and given ad lib for 15 days post busulfan treatment.

### Quantification of tissue findings

Microglia and peripheral monocytes were quantified using imageJ version 1.5s, on flat mount retinas. TUNEL signal was quantified in eye tissue cross-sections stained with DAPI. Retinal ganglion cell quantification was performed manually in Thy1.1^YFP/YFP^ mice and normalized to retinal area. Each analysis was triplicate (n=3). For TUNEL quantification, the area of interest was marked using the freehand selection tool according to DAPI boundaries. The enclosed area was measured (mm^2^) and the image was decomposed to red and blue channels. The red channel that corresponded to TUNEL signal was selected and converted to binary data. The positive pixels (bright pixels) were quantified, the outcome was normalized according to the total sampled area, and the resulting number was plotted.

### Quantification of CX3CR1^+^ cell morphology using grid analysis

The grid-crossed point analysis was used to quantify CX3CR1^+^ cells ramification, per published guidelines (34–36). Briefly, the “grid function” in ImageJ version 1.5s was used to perform the analysis. The grid area per point was adjusted to 625 μm^2^ and the grid was centered on the image. Microglia cells and their processes were traced, and the number of grids each cell occupied was recorded. Frequency plots were generated for each group and statistical analysis was performed to assess differences in frequency distribution between groups. For multiple group comparison, one-way analysis of variance was used with post hoc Tukey’s test. For two group comparison two-sample t-test of normal distribution with unequal variances was performed.

### Retinal immune response analysis with RT^2^ Profiler PCR Array for mouse inflammatory cytokines and receptors

The role of microglia depletion in retinal inflammation was investigated using gene expression analysis. Four groups (3 retinas/group) were analyzed: 1) Control naive C57BL/6J mice (CTRL), 2) Microglia depleted C57BL/6J mice placed on PLX5622 diet for 3 weeks (PLX), 3) C57BL/6J mice that received corneal alkali injury 3 days prior to analysis (Burn), 4) C57BL/6J mice placed on PLX5622 for three weeks followed by corneal alkali injury 3 days prior to analysis (Burn PLX). The analysis required, eye enucleation and surgical dissection of the retinas, tissue homogenization and RNA extraction using RNeasy Mini Kit (74106; Qiagen, Valencia, CA), RNA quantification and normalization using the nanodrop spectrophotometer (Nanodrop 2000; Thermo Fisher Scientific, Waltham, MA), and cDNA synthesis using superscript III (18080-044; Invitrogen, Carlsbad, CA). One microliter of cDNA was loaded in each of the reaction wells of the RT2 Profiler PCR Array Mouse Inflammatory Cytokines & Receptors Kit (PAMM_011Z; Qiagen, San Diego, CA). The assay was performed per manufacturer’s instruction, and data were analyzed by using Qiagen’s data analysis web portal.

### Microglia depletion

Microglia depletion was perform using PLX5622, a CSF1R inhibitor. The drug was provided by Plexxikon Inc. (Berkeley, CA, USA) and formulated in AIN-76A standard chow by Research Diets Inc. (New Brunswick, NJ, USA). A dose of 1200 ppm was given to mice for 3 weeks prior to analysis.

### Statistical analysis

Results were analyzed with the statistical package of social sciences (SPSS) Version 17.0 (Statistical Package for the Social Sciences Inc., Chicago, IL, USA). The normality of continuous variables was assessed using the Kolmogorov-Smirnov test. Quantitative variables were expressed as mean ± standard error of mean (SEM) and qualitative variables were expressed as frequencies and percentages. The Mann-Whitney test was used to assess differences between groups. All tests were two-tailed, and statistical significance was determined at p < 0.05. The independent student t-test was used to compare means between two groups, and pairwise t-test to compare changes within the same group. Analysis of variance (ANOVA) was used for comparisons of multiple groups. Alpha level correction was applied, as appropriate, for multiple comparisons.

## Results

### Microglia depletion exacerbates neuroretinal tissue damage following ocular surface burn injury

We previously showed that corneal injury led to prompt infiltration of peripheral CX3CR1^+^ CCR2^+^ monocytes into the retina and subsequent inflammation that resulted in retinal cell loss (28, 32, 33). In order to assess the role of microglia in this neurodegenerative process we assessed the damage to the retina the setting of microglia depletion by CSF1R inhibitor. CX3CR1^+/EGFP^ and Thy1.1^YFP/YFP^ mice we used to assess the results.

Microglia depletion caused a significant increase in retinal cell apoptosis within 24 hours of the injury, (p<0.05) as compared to injured mice with intact microglia population **(Fig. 1 A-D)**. The absence of microglia in the retina during injury resulted in an increased retinal ganglion cell (RGC) death (P<0.05) and thinning of the nerve fiber layer tissue **(Fig. 1 E-L)** at 2 months, as compared to injured mice with intact microglia. Furthermore, neuroretinal degeneration in microglia depleted mice was associated with epiretinal membrane formation and increased pigmentation from RPE65^+^ cell migration, consistent with proliferative vitreoretinopathy (PVR) disease **(Fig. 1 M-Q)**. In contrast, injured mice with intact microglia exhibited reduced RPE65^+^ cell migration and decreased formation of epiretinal membranes **(Fig. 1 M, O)**.

**Figure 1.**
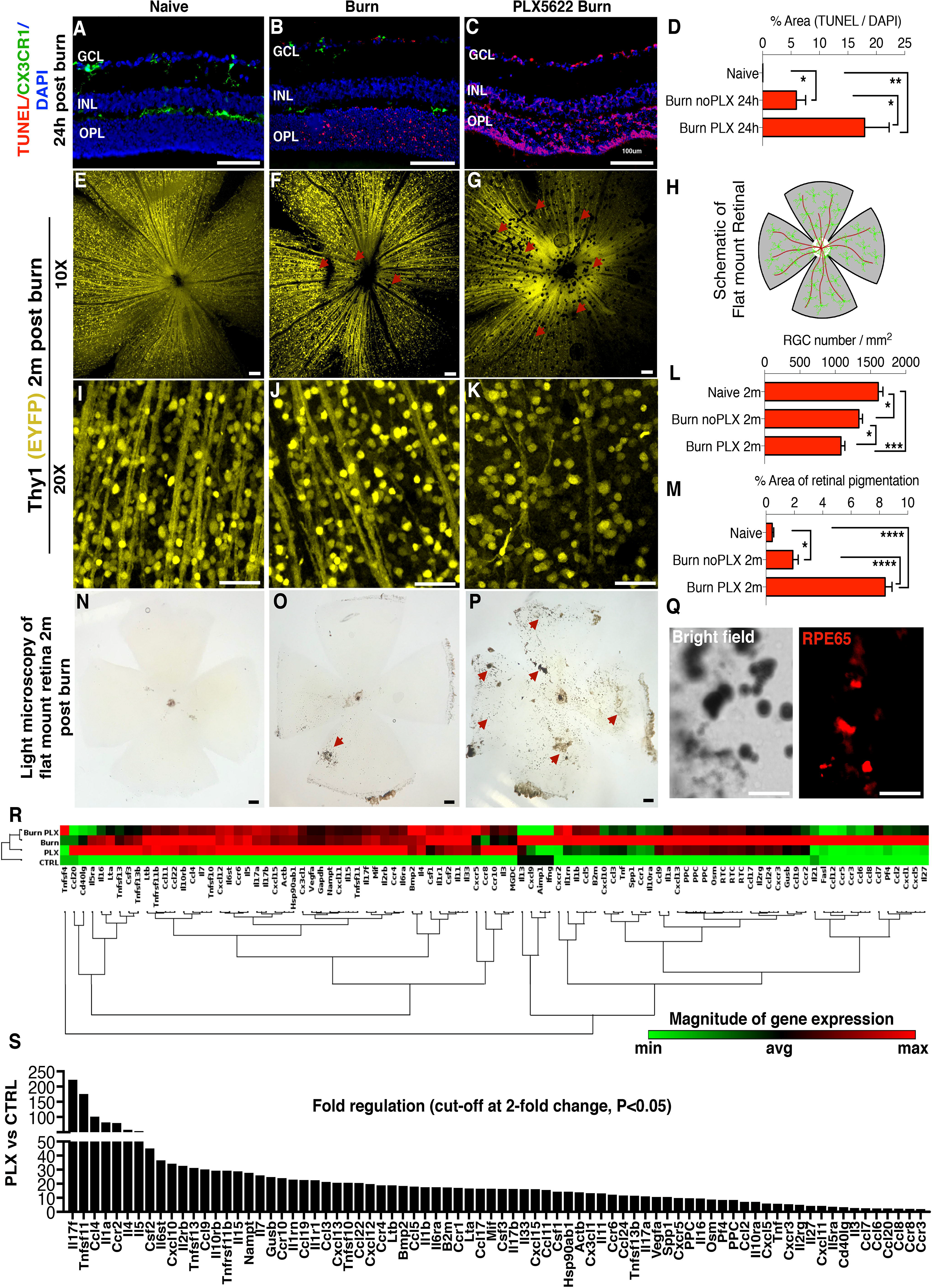
Microglia depletion exacerbates neuroretinal tissue damage after corneal chemical injury. **(A)** TUNEL signal in naive CX3CR1^+/EGFP^ mouse retina. CX3CR1^+/EGFP^ cells are present in the ganglion cell (GCL), inner plexiform/inner nuclear (IPL/INL), and outer plexiform (OPL) layers but TUNEL signal is not present. **(B)** Increase in TUNEL signal in the retina 24 hours after corneal chemical injury. CX3CR1^+/EGFP^ cells are present in all three retinal microglia strata. **(C)** TUNEL signal is significantly elevated 24 hours after ocular injury in microglia depleted mice. **(D)** Quantification of TUNEL signal in the retina 24 hours after the injury. **(E-G)** Images of Thy1 EYFP mice show reduction in retinal nerve fiber layer (RNFL) (red arrows) and **(I-K)** retinal ganglion cells (RGC) 2 months after the ocular injury. However, microglia depleted mice exhibit significantly more RNFL(red arrows) and RGC loss compared to injured mice with intact microglia. **(L)** Quantification of retinal ganglion cell number 2 months after the injury. (**H**) Schematic representation of the flat mount retina. **(M)** Quantification of the pigmented area shows marked increase in microglia depleted mice. **(N)** Light microscopy of naive flat mount, **(O)** 2 months after ocular injury, and **(P)** 2 months after injury in microglia depleted mice. **(N-P)** Ocular injury leads to pigmentation of the inner retina **(P)** (red arrows) which is exacerbated in microglia depleted mice (red arrows). **(Q)** Pigmented cells RPE65^+^ as shown with immunolocalization of flat mount retinas. **(R)** Gene expression analysis of 84 major genes involved in inflammation using RT^2^ Profiler PCR Array of cytokines and receptors in retinas of Naïve C57BL/6 **(CTRL)**, microglia depleted **(PLX)**, corneal burned **(Burn)**, and corneal burned with microglia depletion mice **(Burn PLX)**. Non-supervised hierarchical clustering of the entire dataset with heat map and dendrograms indicating co-regulated genes. Elevation of various inflammatory genes as compared to **CTRL**. **(S)** Microglia depletion leads to significant upregulation of a wide-range of inflammatory genes. Cut-off at 2-fold, P<0.05. Each group n=3 pooled retinas. **(A-Q)** n=5 per group. GCL: ganglion cell layer, IPL/INL: inner plexiform/inner nuclear layer, OPL: outer plexiform layer. **(A-C, N-P)** Scale bar: 100μm, **(E-G, I-K, Q)** Scale bar: 50μm. Multiple comparisons using Tukey’s method \**P*<0.05, \*\**P*<0.01, \*\*\**P*<0.001, \*\*\*\**P*<0.0001.

### Microglia depletion promotes retinal inflammation

We previously showed that microglia are key regulators of peripheral monocyte infiltration into the retina (24). In order to assess the effect of microglial depletion/elimination on the ocular immune response, the expression of 84 major genes involved in inflammation were analyzed in retinas of i) naive **(CTRL)**, ii) microglia depleted **(PLX)**, iii) ocular burn **(Burn)**, and iv) microglia depleted with ocular burn **(Burn PLX)** mice, **(Fig. 1 R)**.

All groups exhibited elevated inflammatory response compared to the **CTLR** group. Genes that were highly (>40-fold) upregulated in all three groups **(PLX, Burn, Burn PLX)** were: *Il-17f, Tnfsf11, Ccr2, Ccl4, Il1a, Il4, Csf2*, and *Il5*. Specific genes highly (>40-fold) upregulated in the **PLX group** were: *Il17f* (220-fold), *Tnfsf11* (175-fold), *Ccl4* (101-fold), *Il1a* (82-fold), *Ccr2* (80-fold), *Il4* (58-fold), *Il5* (53-fold), and *Csf2* (45-fold), among other genes (**Fig. 1 S)**. Specific genes highly (>40-fold) upregulated in the **Burn group** were: *Ccl2* (323-fold), *Ccl6* (296-fold), *Cxcl5* (293-fold), *Il17f* (186-fold), *Ccr2* (155-fold), *Ccl7* (137-fold), *Ccl12* (137-fold), *Tnfsf11* (124-fold), *Ccl4* (113-fold), *Il1a* (108-fold), *Ccr1* (84-fold), *Cxcl10* (84-fold), *Pf4* (79-fold), *Ccr3* (68-fold), *Il27* (64-fold), *Ccl5* (57-fold), *Il1rn* (56-fold), *Csf2* (55-fold), *Il4* (55-fold), Ccl8 (54-fold), *Il5* (52-fold), *B2m* (51-fold), *Ccr5* (50-fold), *Ccl3* (49-fold), *Il1b* (49-fold), *Ccl9* (46-fold) **(Sup. Fig. 1 A)**. Specific genes highly (>40-fold) upregulated in the **Burn PLX group** were: *Il17f* (146-fold), *Cxcl5* (128-fold), *Ccl2* (125-fold), *Tnfsf11* (106-fold), *Ccl4* (84-fold), *Ccl7* (80-fold), *Il4* (63-fold), *Ccr2* (58-fold), *Il1rn* (58-fold), *Csf2* (54-fold), *Il1a* (50-fold), *Il5* (46-fold), *Ccl5* (42-fold) **(Sup. Fig. 1 B)**.

Although many genes exhibited upregulation after burn, depletion of microglia decreases the expression of *Cxcr5*, *Ccl12*, *Ccr5*, *Fasl*, *Ccl6*, and *Ccl8* genes in burned eyes **(Sup. Fig 1 C)**. The following genes were upregulated in both **PLX** and **Burn groups:** *Il-17f, Tnfsf11, Ccr2, Ccl4, Il1a, Il4*, and *Csf2*, while the following genes were upregulated in both the **Burn** and **Burn PLX** groups: *Ccl2*, *Cxcl5*, *Il-17f*, *Ccr2*, *Ccl7*, *Tnfsf11*, *Ccl4*, *Il1a*, *Il4*, *Ccl5*, *Il1rn*, *Csf2*, *Il5*. All above genes were hierarchically arranged according to their expression. Only genes upregulated (>40-fold) were included. The complete range of upregulated genes is shown in **Fig. 1 S** and **Sup. Fig. 1**.

### Microglia depletion causes abnormal engraftment of peripheral CX3CR1^+^ cells in the retina

We previously showed that peripheral CX3CR1^+^ CCR2^+^ monocytes that infiltrate the retina after chemical ocular injury engraft permanently into the tissue and cause neuroglia remodeling (24). These monocytes, although peripheral, were shown to migrate into the three distinct microglia strata, differentiated to ramified cells, and adopted a morphology that resembled microglia. A unique population of peripheral CSF1R^−negative^ CX3CR1^+^ MHC-II^+^ cells was identified within the engrafted cells which contributed remarkably in neuroinflammation and damage to the retinal tissue (24). In order to evaluate the role of native microglia in the process of peripheral monocyte engraftment and subsequent neuroglia remodeling after ocular injury we performed microglia depletion using CSF1R inhibitor PLX5622 in CX3CR1^+/EGFP^ bone marrow chimeras, followed by chemical ocular injury.

Peripheral CX3CR1^+^ cells did not spontaneously infiltrate into the retina under physiological conditions **(Fig. 2 A, E)**, however, ocular chemical injury in bone marrow transferred mice resulted in infiltration and subsequent engraftment of peripheral CX3CR1^+^ cell into the retina at 2 months after the burn **(Fig. 2 B, F, M)**. Engrafted cells migrated uniformly into all three distinct microglia strata (GCL, IPL/INL, OPL) and became highly ramified at 2 months **(Fig 2 N)**. In contrast, engraftment of peripheral CX3CR1^+^ cell in the setting of microglia depletion results was different, with ameboid and semi-ramified peripheral CX3CR1+ monocytes having larger cell bodies and shorter processes and non-uniform distribution **(Fig. 2 C, G)**, as compared to that of injured eyes with intact microglia population **(Fig. 2 B, F)** or to that of naive CX3CR1^+/EGFP^ controls **(Fig. 2 D, H)**. Nevertheless, peripherally engrafted monocytes still managed to migrate into all three distinct microglia strata (GCL, IPL/INL, OPL) by 2 months after the injury **(Fig. 2 Q)**. The mean cell ramification in the BMT burn group **(Fig. 2 I)** was similar to that of BMT PLX burn group **(Fig. 2 J)**,(P= 0.464) but significantly reduced compared to the control CX3CR1^+/EGFP^ group (P< 0.0001), **(Fig 2. K)**.

**Figure 2.**
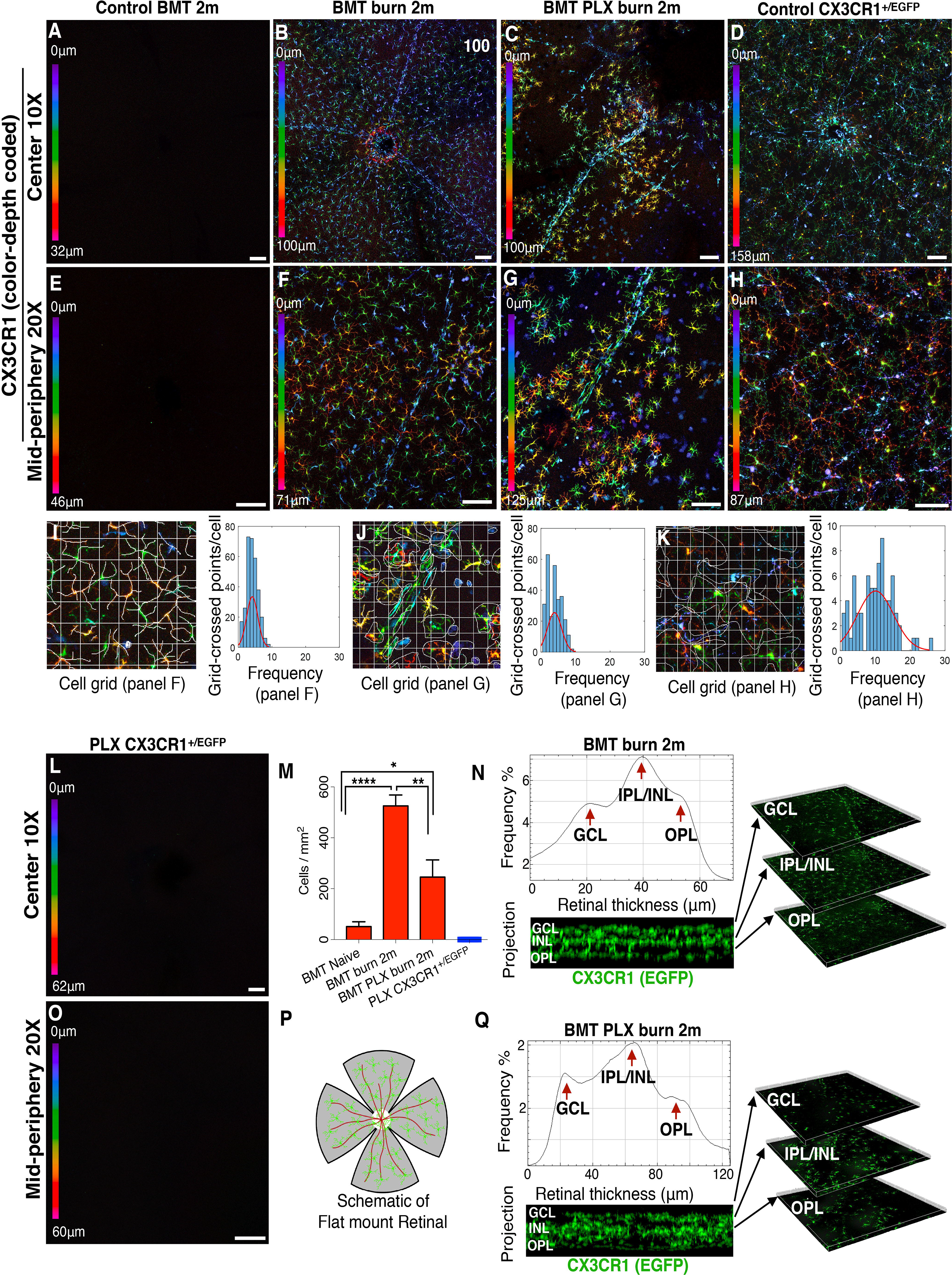
Microglia depletion in injured eyes causes abnormal infiltration and engraftment of peripheral CX3CR1^+^ monocytes into the retina. **(A, E)** Peripheral monocytes do not infiltrate into the retina after busulfan myelodepletion and bone marrow transfer of CX3CR1^+/EGFP^ cells. **(B, F)** Ocular injury causes infiltration of peripheral monocytes in the retina and subsequent engraftment. These cells migrate homogeneously in the retina tissue at 2 months. **(C, G)** Same type of injury in microglia depleted mice results in repopulation of the retina by peripheral CX3CR1^+^ cells at 2 months. Repopulating CX3CR1^+^ cells do not form a homogeneous distribution of cells, but rather form aggregates of cells with semi-ramified morphology. **(D, H)** Normal distribution of CX3CR1^+^ cell in the retina of naive CX3CR1^+/EGFP^ mouse. **(I-K)** Quantification of cell ramification using grid analysis. Representation of the data using frequency plot. Cell ramification in the BMT burn group was similar to BMT PLX burn group (P= 0.464) but significantly reduced compared to control CX3CR1^+/EGFP^ group (P< 0.0001). **(L, O)** Confocal microscopy of retinal flat mounts of CX3CR1^+/EGFP^ mice after 3 weeks of continuous PLX5622 treatment showing complete depletion of CX3CR1^+^ cells from the **(L)** central and **(O)** peripheral retina. **(M)** PLX5622 treatment leads to significant reduction of the number of peripheral CX3CR1^+^ monocytes that engraft into the retina 2 months after the injury. **(N, Q)** Engrafted peripheral monocytes migrate in all 3 distinct microglia strata (GCL: ganglion cell layer, IPL/INL: inner plexiform/inner nuclear layer, OPL: outer plexiform layer) 2 months after ocular injury in mice with intact microglia system as well as in mice with depleted microglia. **(P)** Schematic representation of flat mount retina used for confocal microscopy. n=5 per group. GCL: ganglion cell layer, IPL/INL: inner plexiform/inner nuclear layer, OPL: outer plexiform layer, BMT: bone marrow transfer. **(A-D, L)** Scale bar: 100μm, **(E-H, O)** Scale bar: 50μm. Multiple comparisons using Tukey’s method \**P<0.05*, \*\**P<0.01*, \*\*\*\**P<0.0001*.

Although PLX5622 treatment completely depleted retinal CX3CR1^+/EGFP^ microglia **(Fig. 2 L, O)**, replenishment of these cells occurred from peripheral monocyte infiltration (24). Likewise, peripheral monocytes infiltrated the retina of non-depleted mice after burn, however, the number of peripheral CX3CR1^+^ infiltrates at 2 months was markedly reduced in microglia depleted mice as compared to non-depleted **(Fig. 2 M)**. A schematic representation of flat mounted retina is shown **(Fig. 2 P)**.

### Ocular hypertension, penetrating corneal injury, but <u>not</u> optic nerve crush, cause peripheral monocyte engraftment into the retina

Our previous findings showed that corneal chemical injury caused peripheral CX3CR1^+^ monocyte infiltration and engraftment into the retina and subsequent pigmentation of the inner retina. In order to assess if other types of ocular injuries also induce the same changes, we employed three clinically relevant ocular injuries models: a) acute ocular hypertension that recapitulates acute angle closure glaucoma, b) penetrating corneal injury/surgical trauma, and c) optic nerve crush that recapitulates traumatic optic nerve injury.

#### Ocular hypertension injury

Firstly we confirmed that peripheral monocytes do not enter into the retina without injury **(Fig. 3 A, D)**. Acute ocular hypertension caused significant infiltration and subsequent engraftment of peripheral CX3CR1^+^ monocytes into the retina within 2 months **(Fig. 3 B, G, E)**. Engrafted peripheral monocytes migrated in all three distinct microglia strata **(Fig. 3 C)** and transformed to dendritiform cells **(Fig. 3 E)**. Although peripheral monocytes that infiltrated the retina after OHT injury adopted a ramified morphology **(Fig 3. F)**, the extend of ramification remained significantly lower (*P*<*0.0001*) compared to resident microglia **(Fig. 2 K)**. Aside from RGC loss, acute ocular hypertension caused significant damage to the retina, as evident by RPE65^+^ cell migration and pigmentation of the inner retina **(Fig. 3 H, I, J)**. A schematic representation of flat mounted retina is shown **(Fig. 3 K)**.

**Figure 3.**
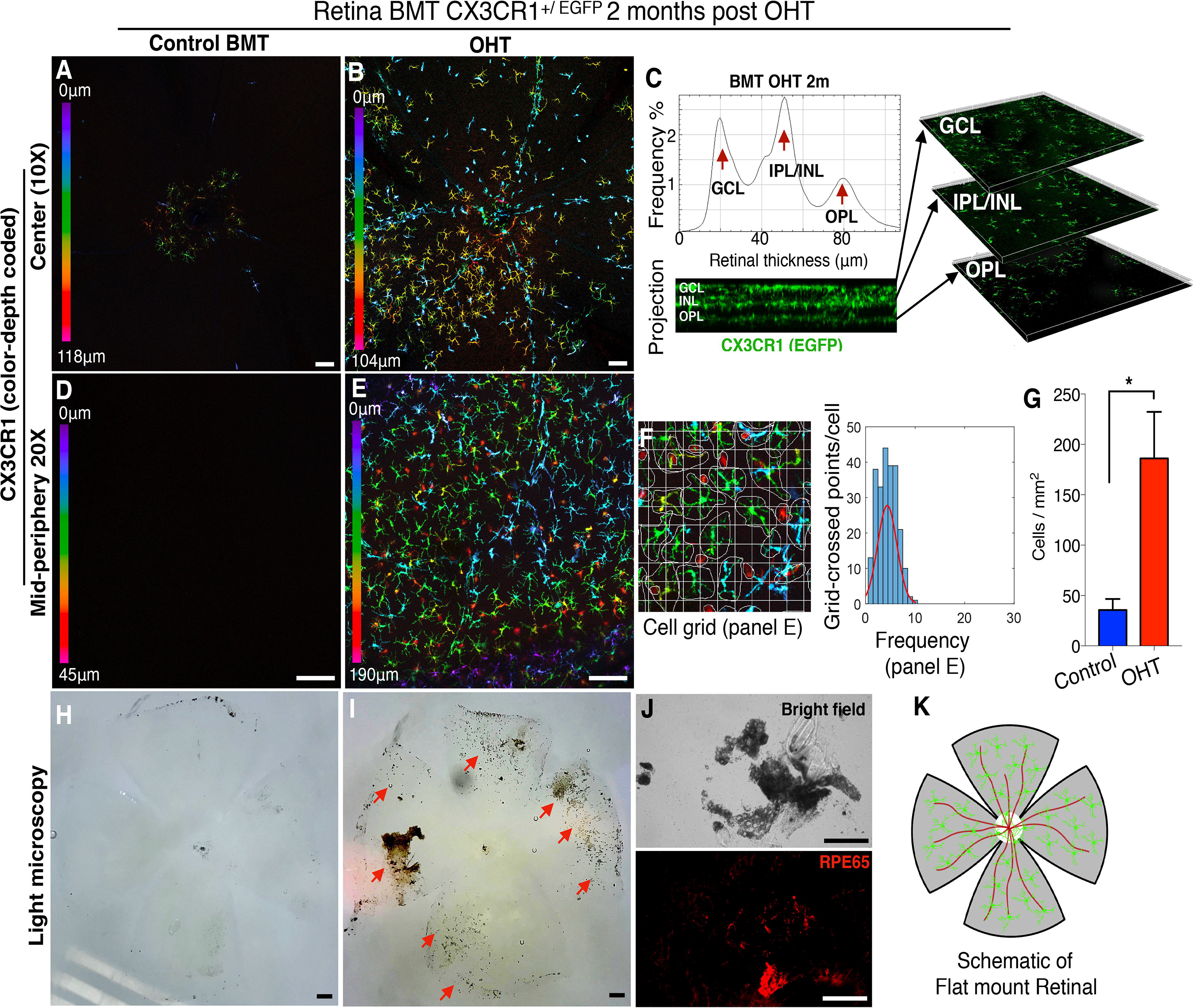
Acute ocular hypertension causes peripheral CX3CR1^+^ infiltration and engraftment into the retina. Ocular hypertension (OHT) in BMT CX3CR1^+/EGFP^ mice using anterior chamber cannulation (intraocular pressure elevation to 100 mmHg for 45 minutes). **(A, D)** In control mice CX3CR1^+^ cells are present in the retina 2 months after BMT. **(B, G, E)** OHT causes significant infiltration (P<0.05) of peripheral CX3CR1^+^ monocytes into the retina and subsequent differentiation to ramified cells within 2 months. **(C)** Infiltrated peripheral CX3CR1^+^ monocytes migrate in all three distinct microglia strata (ganglion cell, inner nuclear cell, and outer plexiform cell layers) 2 months after the injury and engraft into the tissue. **(F)** Peripheral monocytes that infiltrate the retina after OHT injury adopt a ramified morphology but remain less ramified (P<0.0001) compared to resident (native) microglia. **(H)** In control eyes, the inner retina shows no evidence of secondary pigmentation. **(I, J)** Two months after the injury, the inner retina becomes significantly pigmented (red arrows) with RPE65^+^ cells. **(K)** Schematic representation of flat mount retina used for confocal microscopy. BMT: bone marrow transfer, OHT: ocular hypertension. n=3 per group. **(A, B, D, E, H, I, J)** Scale bar: 100μm. Student t-test \**P*<0.05.

To confirm that the observed monocyte response after acute ocular hypertension is due to pressure elevation and not to injury by the cannulation, we performed mock experiments in which the cannula was placed in the anterior chamber for 45 minutes but the intraocular pressure was not elevated. Mock induction did not cause peripheral CX3CR1^+^ cell infiltration into the retina, as observed with confocal microscopy at 2 months after the procedure **(Sup. Fig. 1 A-D)**.

Likewise, penetrating corneal injury using full thickness corneal suture **(Fig. 4 A)** caused peripheral CX3CR1^+^ cell engraftment into the retina (Fig. 4 B, C, E, F). These cells uniformly occupied all three distinct microglia strata and adopted a ramified morphology **(Fig. 4 C, F, G)** as compared to eyes without injury **(Fig. 4 B, E)**. Peripheral monocytes that infiltrated the retina after penetrating ocular injury were more ramified compared to infiltrating peripheral monocytes of control BMT mice (P<0.0001), **(Fig 4. H, F)** but also less ramified as compare to resident microglia (P<0.0001), **(Fig. 2 K)**. In addition, penetrating corneal injury caused significant damage to the retina, as evident by RPE65^+^ cell migration and pigmentation of the retina **(Fig. 4 J-L)**. A schematic representation of flat mounted retina is shown **(Fig. 4 M)**.

**Figure 4.**
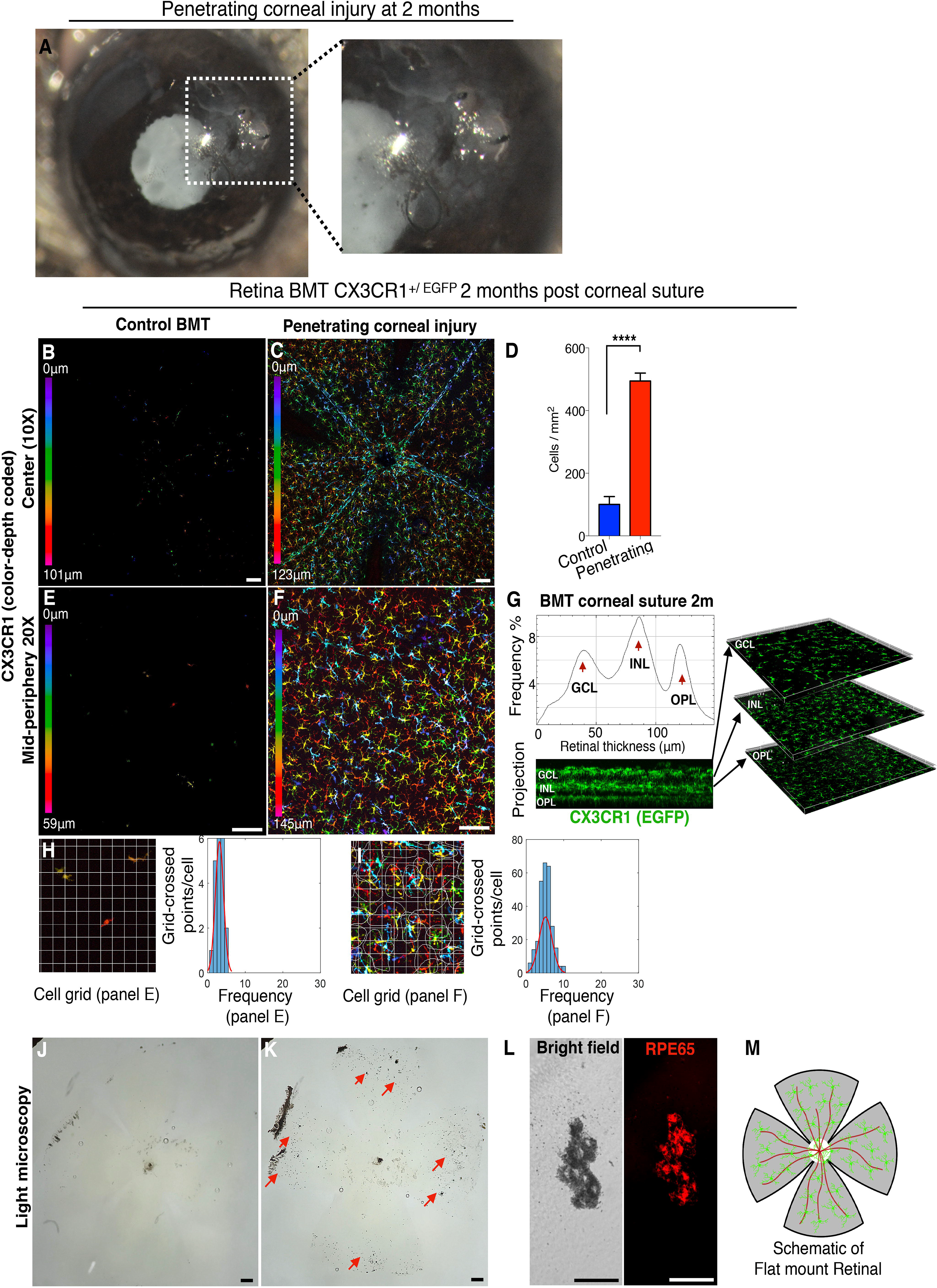
Penetrating corneal injury leads to peripheral CX3CR1^+^ infiltration and engraftment into the retina. **(A)** Penetrating cornea injury performed by full-thickness placement of 11-0 vycril suture in the cornea. **(B, E)** CX3CR1^+/EGFP^ bone marrow chimera model shows no peripheral monocyte infiltration in the absence of a suture. **(C, D, F)** Penetrating corneal injury causes engraftment of peripheral CX3CR1^+^ monocytes into the retina with a ramified appearance at 2 months. **(G)** Engrafted peripheral CX3CR1^+^ monocytes migrate into the three distinct microglia strata (ganglion cell, inner nuclear cell, and outer plexiform cell layers) 2 months after the injury. **(H, I)** Peripheral monocytes that infiltrate the retina after penetrating ocular injury are more ramified than in control BMT mice (P<0.0001) and less ramified compared to resident (native) microglia (P<0.0001). **(J-L)** Penetrating corneal injury causes pigmentation of the inner retina (red arrows) by RPE65^+^ cells. **(M)** Schematic representation of flat mount retina used for confocal microscopy. BMT: bone marrow transfer, OHT: ocular hypertension. n=3 per group. **(B, C, E, F, J, K)** Scale bar: 100μm. **(L)** Scale bar: 50μm. Student t-test \*\*\*\**P<0.0001*.

#### Optic nerve crush injury

In contrast, optic nerve crush-mediated RGC loss was not associated with peripheral CX3CR1^+^ monocyte infiltration into the retina **(Fig. 5 A, B, E, F, I)**, nor PVR **(Fig. 5 C, G)**, despite the fact that this injury is known to cause dramatic loss of RGCs and thinning of the nerve fiber layer within 2 months (37, 38). Also, optic nerve crush did not cause RPE65^+^ cell migration nor pigmentation of the inner retina **(Fig. 5 C)**. Interestingly, peripheral monocytes that infiltrated the injured optic nerve were separable into CCR2^+^ CX3CR1^−negative^ cells present at the site of the injury, and CCR2^−negative^ CX3CR1^+^ cells seen at the adjacent uninjured tissue **(Fig. 5 D & Sup. Video 1)**. We have previously shown that peripheral CX3CR1^+^ and CCR2^+^ cells physiologically populate the optic nerve **(Fig. 5 H)** but do not infiltrate the retina (28). Likewise, optic nerve crush caused marked increase in CX3CR1^+^ and CCR2^+^ cells in either sides of the optic nerve crush **(Fig. 5 D)**, suggesting that the injury site is not a barrier to the migration of peripheral monocytes into the retina. A schematic representation of flat mounted retina is shown **(Fig. 5 J)**.

**Figure 5.**
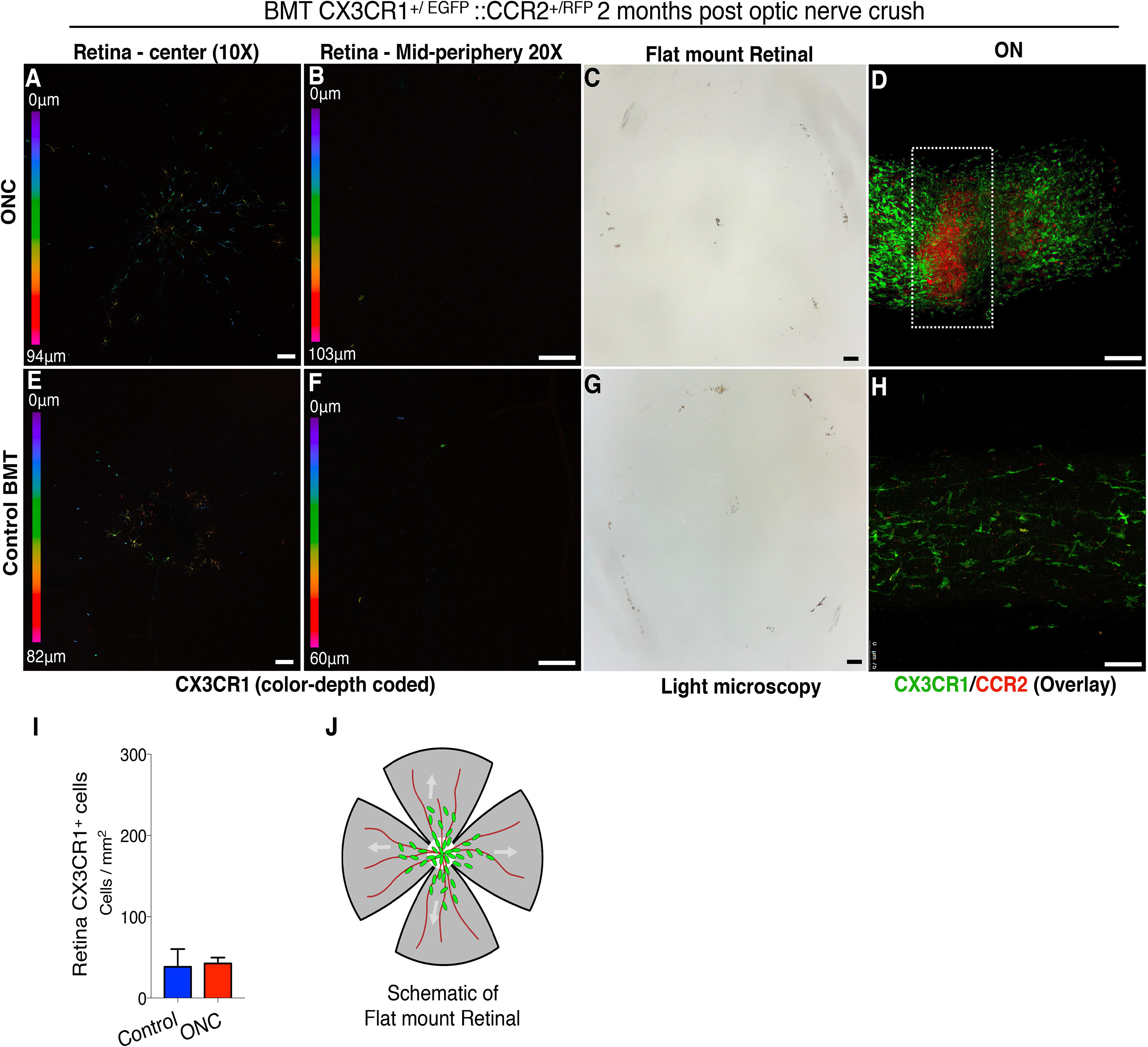
Optic nerve crush does <u>not</u> cause peripheral CX3CR1^+^ infiltration and engraftment into the retinal. Optic nerve crush (ONC) injury in CX3CR1^+/EGFP^::CCR2^+/RFP^ bone marrow chimeras. **(A, B, E**, ONC does not cause peripheral CX3CR1^+^ CCR2^+^ cell infiltration or engraftment into the retina, as assessed at 2 months or inner retina pigmentation **(C, G)**. **(D, H)** Flat mount confocal microscopy of the optic nerve shows a significant increase in peripheral CCR2^+^ CX3CR1^−negative^ cells at the site of the ONC, and CCR2^−negative^ CX3CR1^+^ cells adjacent to the ONC, in the uninjured tissue. **(I)** No engraftment of peripheral CX3CR+ cells in the retina is seen 2 months after ONC injury. **(J)** Schematic representation of flat mount retina used for confocal microscopy. BMT: bone marrow transfer, ONC: optic nerve crush. n=6. **(A-H)** Scale bar: 100μm. Student t-test \*\*\*\**P*<*0.0001*.

## Discussion

Our study suggests that corneal chemical injury, penetrating injury, and ocular hypertension, but not optic nerve crush, trigger similar neuroglia remodeling and damage to the retina. We show that after ocular injury, peripheral monocytes engraft permanently into the tissue, migrate into the three distinct microglia strata (GCL, IPL/INL, OPL), adopt a ramified morphology, and establish a pro-inflammatory neuroglia phenotype that promotes neurodegeneration (24). These findings are likely important for patients with surgical trauma, such as penetrating keratoplasty or keratoprosthesis surgery, that have been shown to exhibit progressive neurodegeneration, even in the absence of IOP elevation (39). According to our data, neuroglia remodeling through peripheral monocyte engraftment contributes in pathogenesis and progression of retinal degeneration, a hypothesis that is also supported by other recent animal studies (19, 32).

Our data suggest that microglia may regulate the process of peripheral monocyte infiltration into the retina, facilitate their engraftment, and regulate damage to retinal neurons. Microglia depletion during ocular injury exacerbated retinal damage and resulted in abnormal engraftment of peripheral monocytes into the tissue. In contrast, the same injury with microglia intact resulted in reduced retinal damage and a normal morphology of engrafted peripheral monocytes. It appears that microglia provide local cues that facilitate integration of peripheral monocytes into the pool of retinal microglia. Conversely, microglia depletion during injury affects this process and results in abnormal integration. The ability of peripheral monocytes to integrate into the brain microglia pool has been demonstrated during bacterial infection of the CNS, where blood monocytes/macrophages engrafted into the brain and became actively involved in the resolution of tissue damage (40). Likewise, peripheral monocyte engraftment in the brain has been demonstrated in a mouse model of Parkinson’s disease (41) in meningitis (42), in Alzheimer’s disease (43), and during early incubation period of scrapie (44). In this study we show that various ocular injuries, including surgical trauma, may cause a similar process in the retina.

Preferential recruitment and engraftment of peripheral monocytes occurs early in various neuropathologies (24, 44) and is not always the result of blood barrier disruption (19, 43, 44). Microglia depletion was associated with significant upregulation of several major inflammatory genes in the retina, suggesting that microglia elimination leads to loss of retinal homeostasis and increased inflammation **(Fig. 1 S)**. Upregulation of proinflammatory molecules, such as *Il17f*, *Ccr2*, *Tnfα*, *Il1α*, *Csf2*, *Il1β*, *Il4* and *Il6* were seen in our study. Several of them have been shown elsewhere to mediate peripheral monocyte recruitment into the retina and their blockade reduces this infiltration (19, 32). The role of engrafted peripheral monocytes in the CNS, however, remains elusive. Previous studies have shown that peripherally engrafted monocytes contribute to disease clearance of scrapie (44). We recently demonstrated that engraftment of peripheral CX3CR1^+^ CSF1R^−negative^ monocytes into the retina is pathologic and contributes to neuroretinal degeneration (24). Herein, we show that various ocular injuries that do not directly effect the retina may still instigate similar neuroglia remodeling and damage to the neurons through peripheral monocyte engraftment. In this process, microglia retain a protective role that is overshadowed by the engrafted monocytes. Therefore, the functional duality observed for microglia after injury may very well reflect the transcriptional heterogeneity of the various monocytic subtypes that comprise the neuroglia system after it has been remodeled.

Surprisingly, optic nerve crush did not cause peripheral monocyte infiltration into the retina, although peripheral monocytes massively infiltrate the injured optic nerve tissue. As previously demonstrated, RGC loss after optic nerve crush (37, 45) does not appear to depend on peripheral monocyte infiltration, corroborating our findings. Optic nerve crush does not appear to involve retinal microglia activation either, since a recent study showed that microglia are irrelevant for neuronal degeneration and axon regeneration after optic nerve crush injury and therefore their depletion does not affect this process (45). Although optic nerve crush leads to significant RGC loss, for various reasons, the contribution of peripheral monocytes appears insignificant.

The failure of peripheral CX3CR1^+^ monocytes to infiltrate the retina after optic nerve crush was not attributed to the physical barrier caused by the nerve trauma; peripheral CX3CR1^+^ monocytes were present on both sides (proximal and distal) of the injured tissue as well as around the optic nerve head. We previously showed that although peripheral monocytes physiologically populate the optic nerve and become tissue resident macrophages (19), they do not migrate into the retina (19, 24), despite the physical proximity of the two tissues. It appears that peripheral monocytes are differentially regulated in regards to retina and optic nerve migration (19). In this regard, resident retinal microglia appear to have a pivotal role as barriers to this migration (24).

Our results suggest that microglia also regulate RPE65^+^ cell migration into the retina after injury. Although all ocular injuries (penetrating corneal injury, ocular hypertension, and corneal chemical injury) resulted in RPE65^+^ cell migration and pigmentation of the inner retina, microglia depletion significantly increased both findings. These findings are likely clinically significant as RPE cell migration into the retina is known to be a contributor in PVR (46), and PVR is a major contributor to retinal detachment (47). RPE cells that migrate into the retina undergo transformation, deposit extracellular matrix, and form epiretinal membranes (48–51).

Such membranes contain immune cells (47, 52), including microglia and peripheral macrophages (48, 49, 53, 54). Preventing RPE cell migration improves the clinical outcomes after ocular injury (52), and therefore, preserving native microglia may be beneficial. Our study shows that RPE migration starts from the periphery **(Fig. 3 J)**. Indeed, an increased number of pigmented cells were present in the peripheral retina relative to the central retina. Although more work is necessary to understand the role of microglia in RPE regulation, our study provided evidence to suggest that microglia suppress RPE migration and PVR, and that restoration or preservation of the physiological microglial phenotype may be a potential therapeutic target. Another possible therapeutic strategy to protect the retina may be the modulation of microglia activity as opposed to inhibition. Microglia, as well as macrophages can acquire signatures that promote or ameliorate inflammation. For example, reactive microglia may secrete inflammatory mediators, such as colony stimulating factors (M-CSF, GM-CSF), TNF-α and IL-1β, IL-6, but may also secrete neuroprotective factors, such as brain-derived neurotrophic factor (BDNF), glial-derived neurotrophic factor (GDNF), nerve growth factor (NGF), neurotrophin 3 (NT-3), NT-4 (55, 56). The balance between inflammatory/protective expression is what determines the outcome.

Previous studies by Gibbings(57), van Der Laar (58), as well as ours(24), suggest that epigenetic reprogramming of repopulating peripheral monocytes may happen by local tissue cues, such as inflammation. This opens possibilities of modulating microglia environment to support expressions that support the neurons. Thus, altering the signature of these cells to a more protective phenotype could be an imperative therapeutic approach for the future. Epigenetic reprogramming methods that can alter the phenotype of microglia or peripheral monocytes have been recently discussed(24), however, this field requires substantially more work to transition into clinical reality.

In conclusion, our findings suggest that microglia cells are not only key regulators of peripheral monocyte engraftment into the retina after ocular injury, but also suppressors of RPE cell migration into the retina. Microglia depletion leads to significant upregulation of several major inflammatory genes in the retina and loss of retinal homeostasis. Although retinal degeneration occurs in many different injury models, injury to the optic nerve does not result in peripheral monocyte infiltration nor cause RPE65^+^ cell migration. Future studies need to account for changes in retinal microglia phenotype after injury due to peripheral monocyte engraftment, rather than transcriptional changes from native microglia. The latter may also occur, but more likely promotes neuroprotection. From a therapeutic standpoint, reestablishing normal neuroglia phenotype may be possible through selective ablation or reprogramming of peripherally engrafted monocytes while preserving the normal function of resident microglia.

## Acknowledgments

This work was supported by the Boston Keratoprosthesis Research Fund, Massachusetts Eye and Ear, the Eleanor and Miles Shore Fund, the Massachusetts Lions Eye Research Fund, an unrestricted grant to the Department of Ophthalmology, Harvard Medical School, from Research to Prevent Blindness, NY, NY, and NIH National Eye Institute core grant P30EY003790.

---

**Sup. Figure 1.**
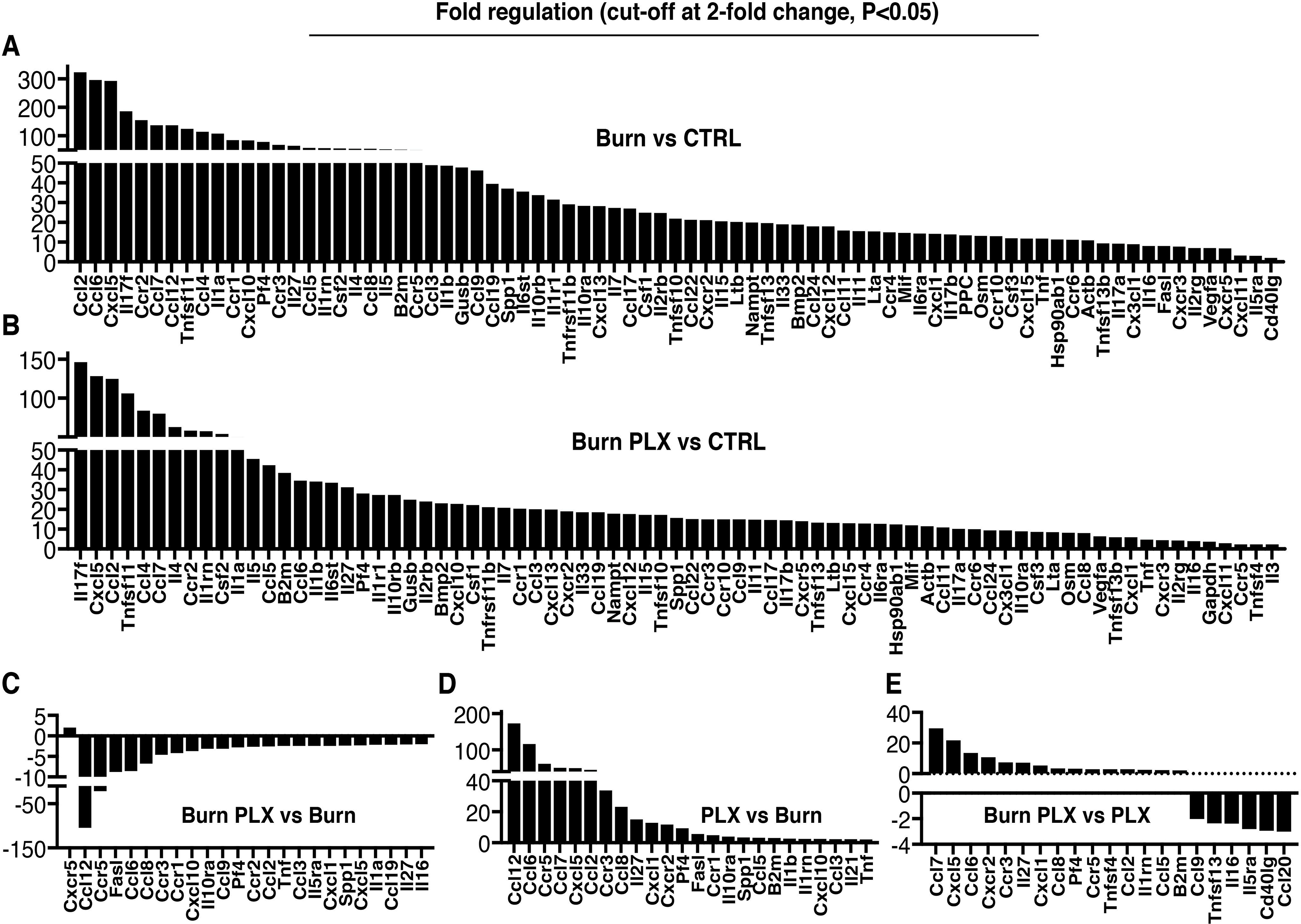
Gene expression changes after microglia depletion. Retinal gene expression analysis of major inflammatory markers using RT2 Profiler PCR Arrays in **1)** control naive C57BL/6 **(CTRL), 2)** microglia depleted C57BL/6 **(PLX), 3)** corneal burned **(Burn)**, and **4)** corneal burned with depleted microglia mice **(Burn PLX)**. Significant gene expression differences between **(A)** Burn and CTRL, **(B)** Bum PLX and CTRL, **(C)** Burn PLX and Burn, **(D)** PLX and Burn, and **(E)** Burn PLX and PLX groups. Significance cut-off limit was set to 2-fold expression change, P<0.05.

**Sup. Figure 2.**
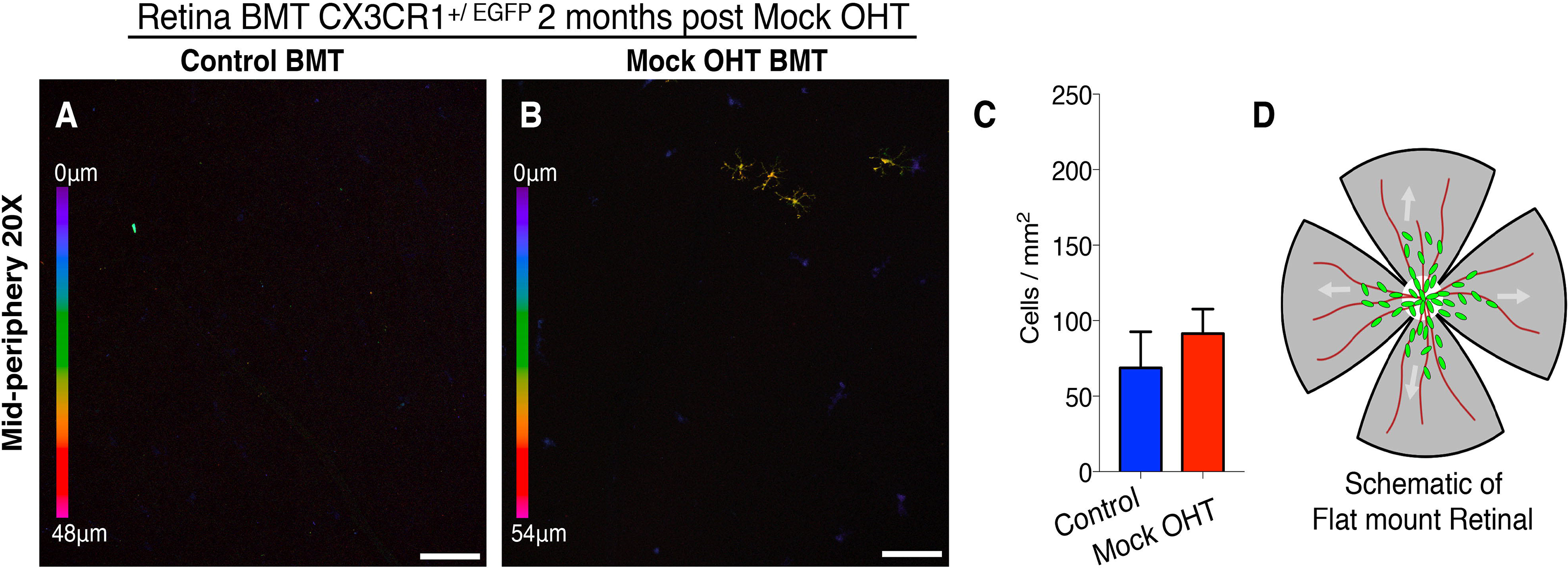
Mock ocular hypertension does not causes peripheral CX3CR1+ infiltration and into the retinal. Mock ocular hypertension (OHT) experiments in BMT CX3CR1^+/EGFP^ mice performed by placing the cannula in the
anterior chamber for 45 minutes without elevating the intraocular pressure. **(A-C)** Mock OHT does not causes peripheral CX3CR1+ monocytes infiltration into the retina at 2 months. **(D)** Schematic representation of flat mount retina used for confocal microscopy. BMT: bone marrow transfer, OHT: ocular hypertension. n=3 per group. **(A, B)** Scale bar: lOOμm. Student t-test.

